## Supplementary Figures for "Spatially Interacting Phosphorylation Sites and Mutations in Cancer"

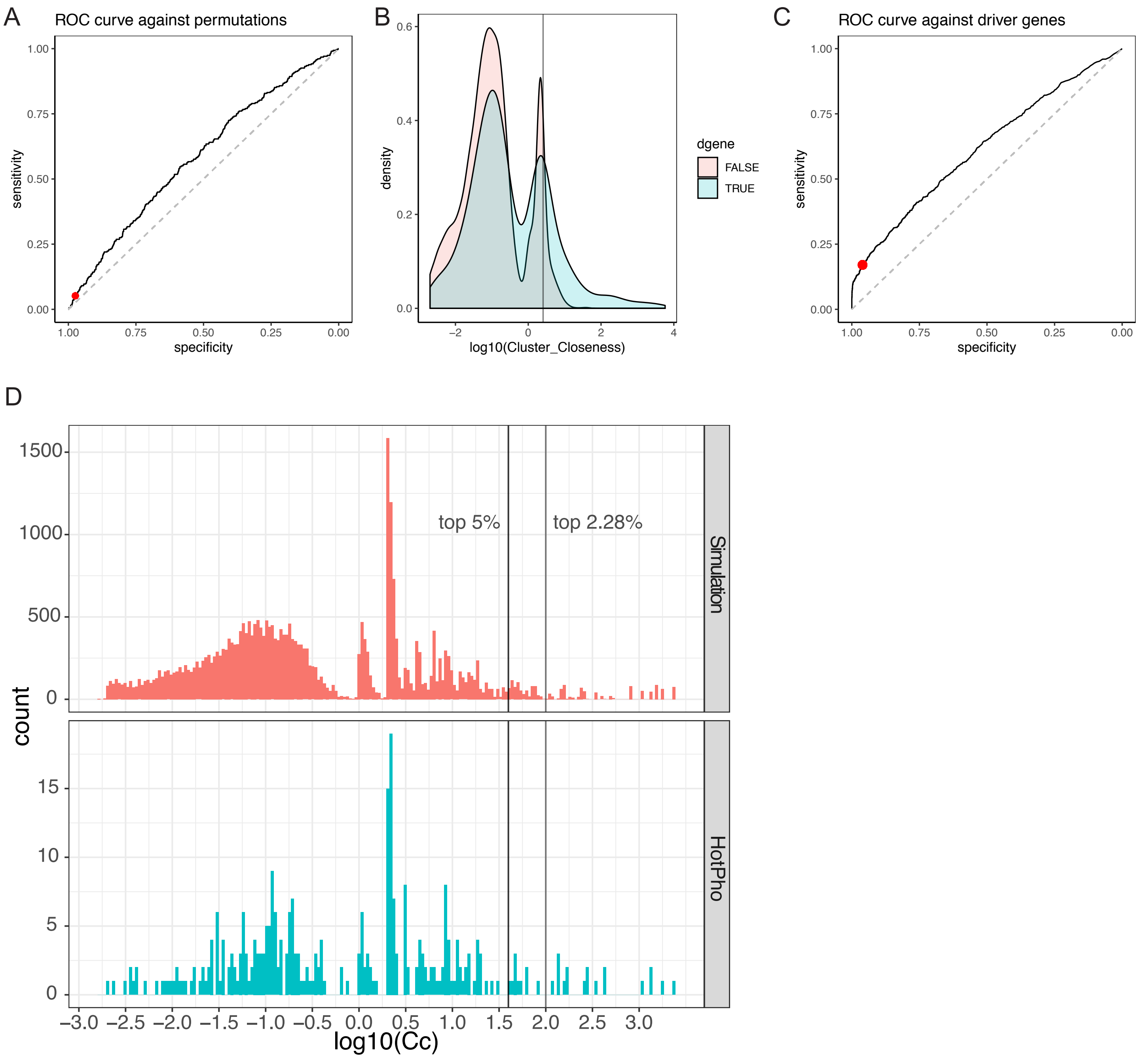


**Supplementary Figure 1.HotPho Benchmarks.** (A) ROC curve analysis of HotPho using bona fide data vs. simulated data of phosphosites. (B) The cluster closeness score density distribution for the 299 driver genes (dgene) vs. other genes. (C) ROC curve analysis of HotPho hybrid clusters in distinguishing clusters involving driver genes and other clusters. For (A) and (C), The red dot indicates the score threshold of top 5% used in the manuscript. (D) A histogram of the cluster-closeness scores in HotPho-generated hybrid clusters using observed (HotPho) vs. simulated data.

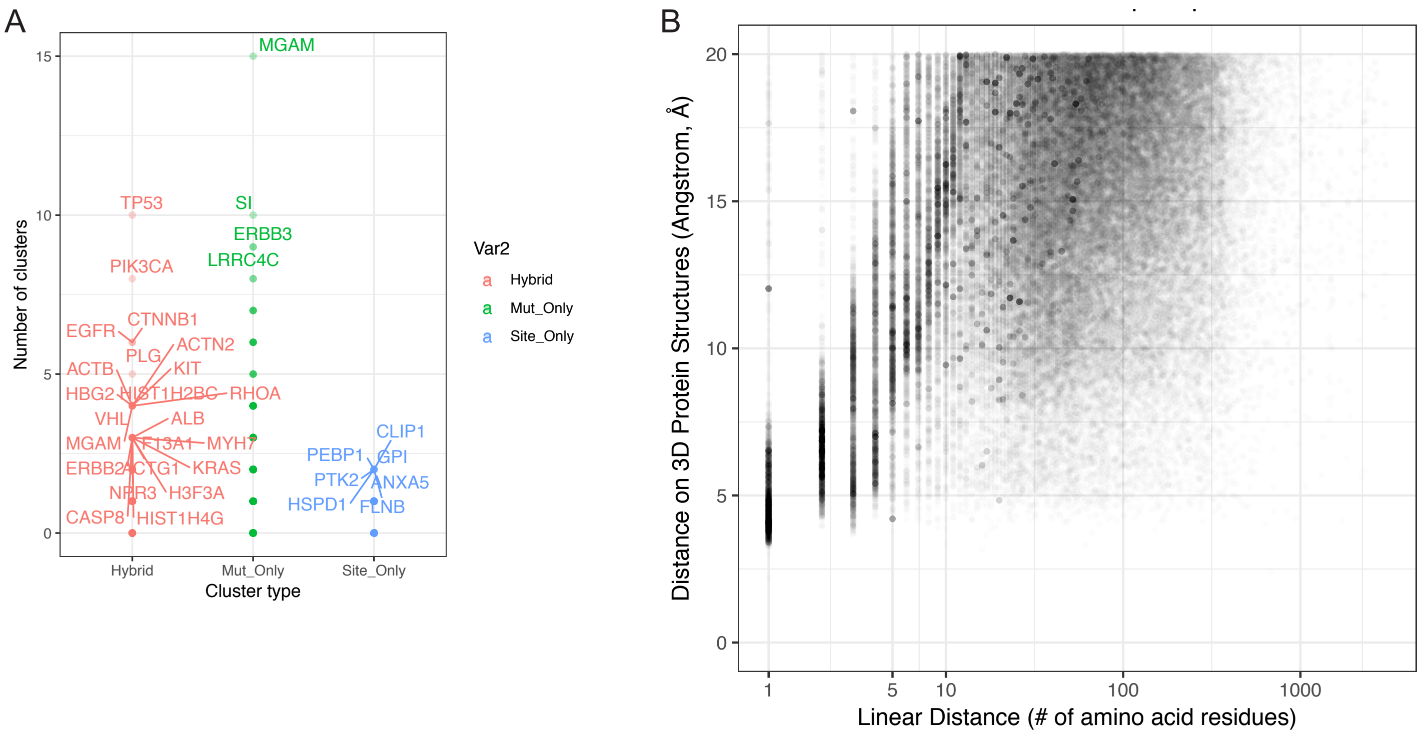


**Supplementary Figure 2.Three-dimensional clustering of phosphosites and mutations.** (A) Number of mutation-only, phosphosite-only, and hybrid clusters detected in each protein; proteins with the highest cluster counts in each categories are highlighted. (B) Linear and 3D Distance between co-clustered mutations and phosphosites identified by HotPho.

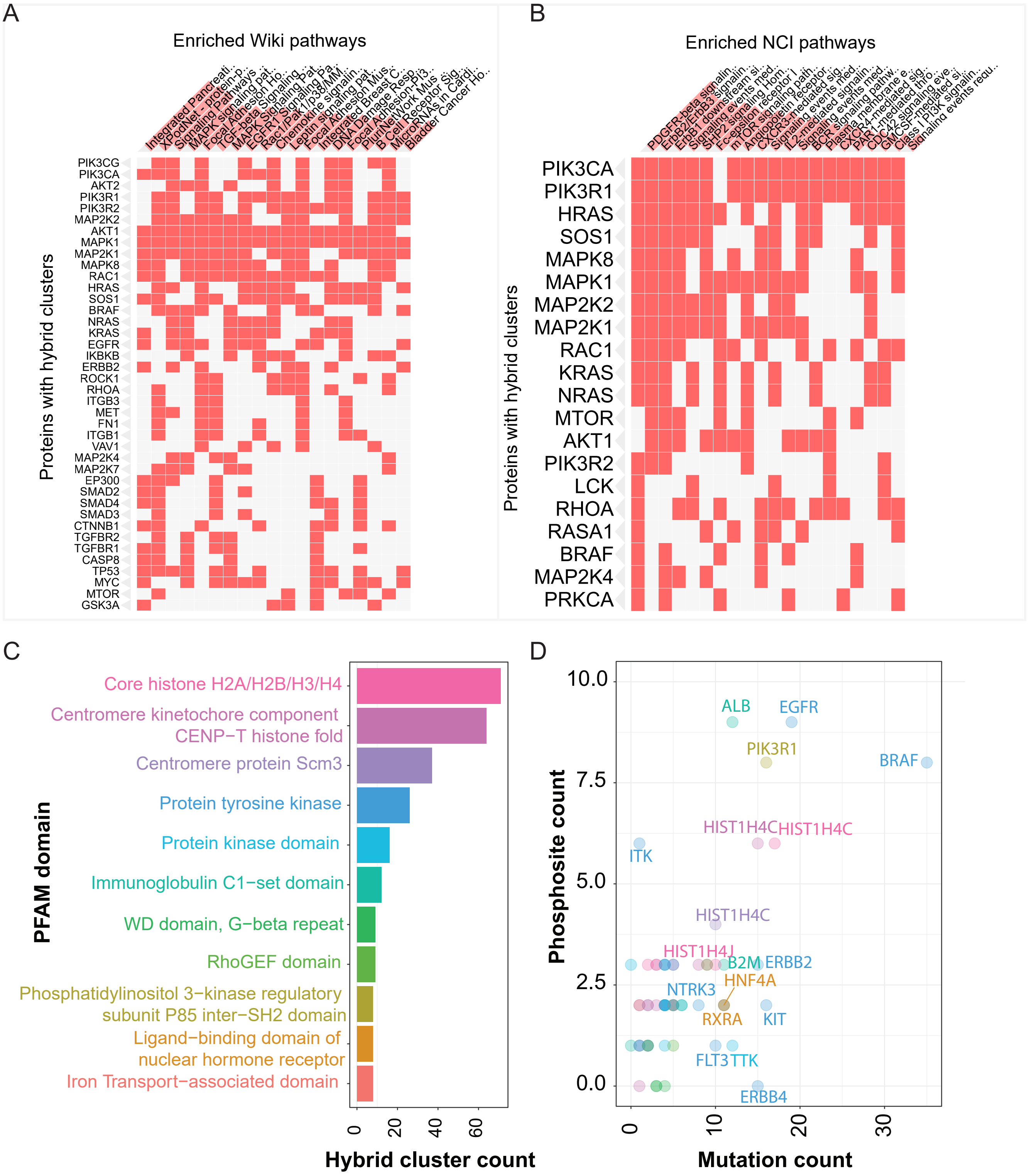


**Supplementary Figure 3. Pathway and domain enrichment analyses of hybrid clusters.** (A) Top WikiPathways and (B) NCI pathways associated with proteins with hybrid clusters identified using HotPho. Each column indicates enriched pathways (Full pathway name in Table S2) and row indicates proteins with hybrid clusters with membership (shown in red blocks) in the specific pathways. (C) Top 11 Pfam domains that each contains more than 7 hybrid clusters. (D) Number of phosphosites and mutations of hybrid clusters overlapping the top 11 Pfam domains. Each dot represents a hybrid cluster and only hybrid clusters with more than 3 co-clustering phosphosites or 9 co-clustering mutations are highlighted with their gene names.

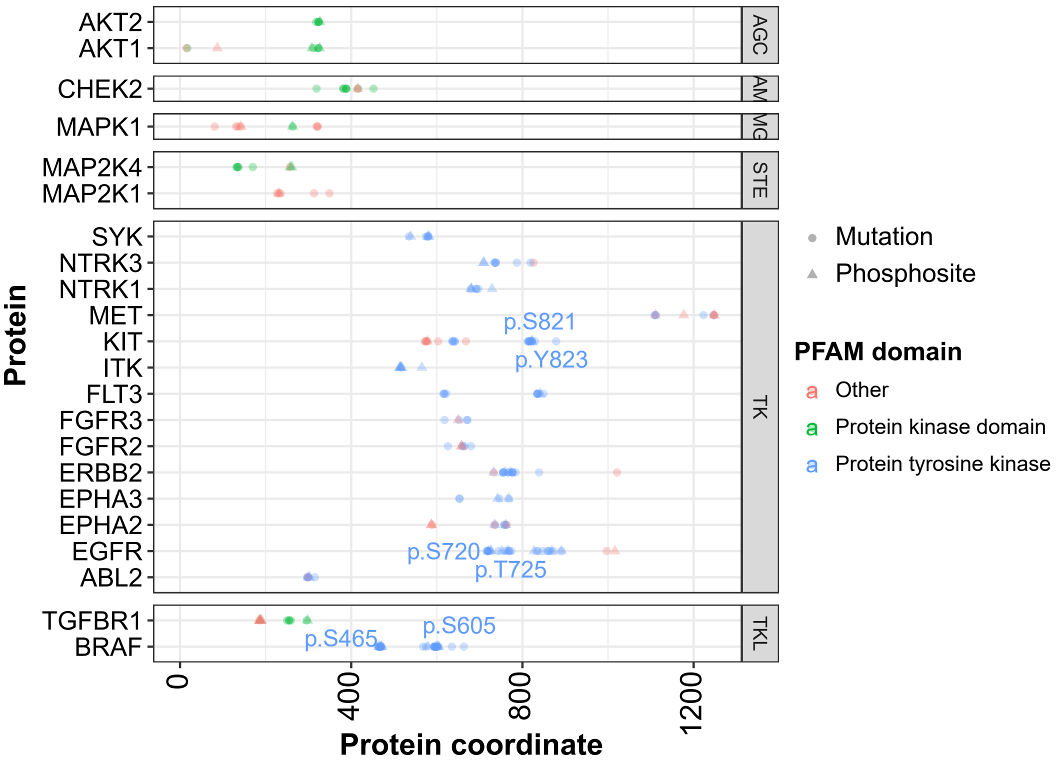


**Supplementary Figure 4.** Co-clustering mutations and phosphosites located in the protein tyrosine kinase and protein kinase domains across kinases of major kinase groups.

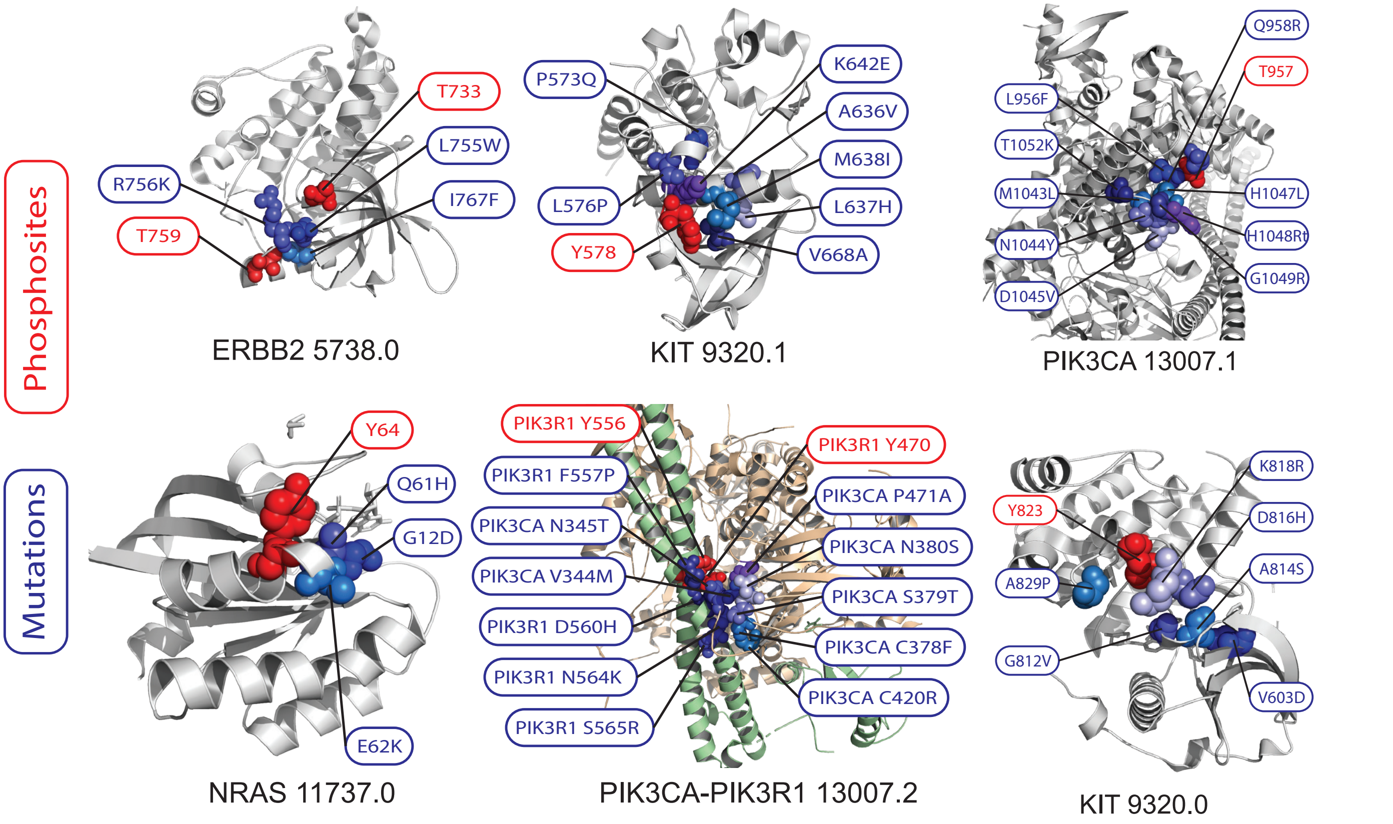


**Supplementary Figure 5.** Selected hybrid clusters with activating or recurrent mutations shown on 3D protein structures obtained through PDB. Mutations are colored in shades of blue and phosphosites are colored in shades of red.

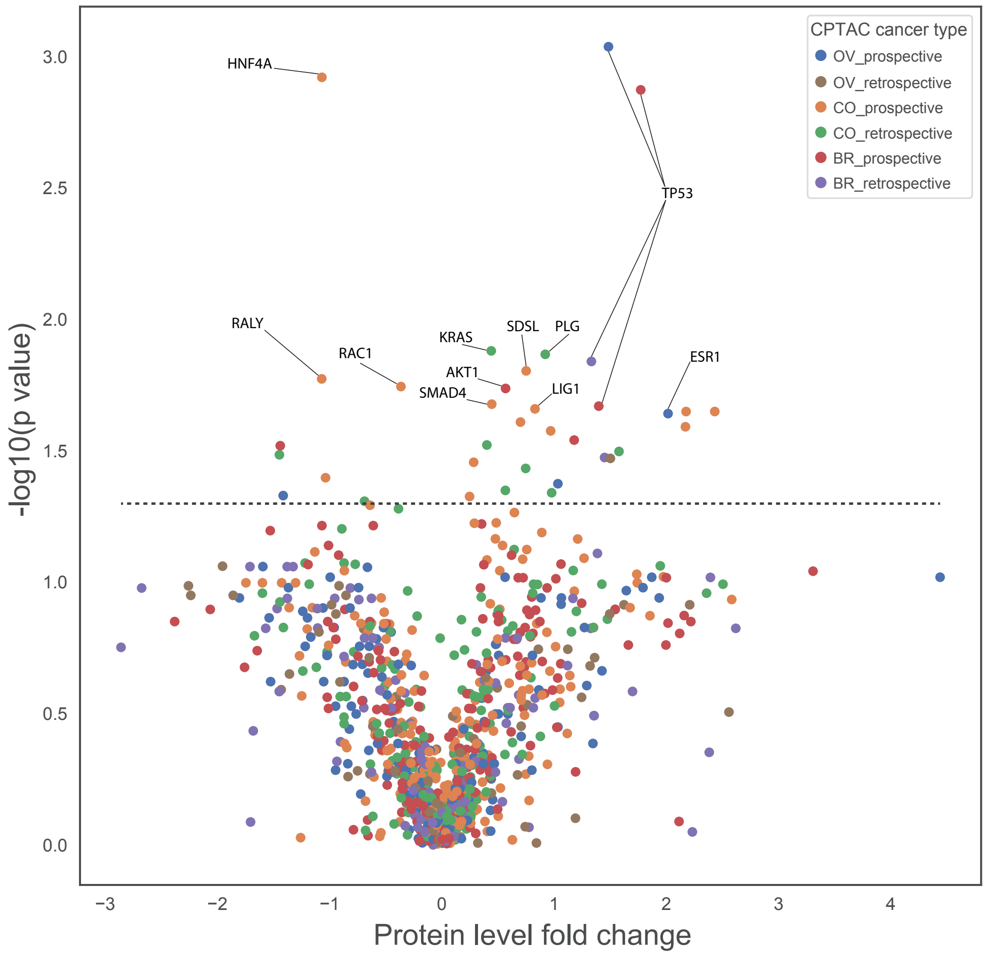


**Supplementary Figure 6.** Volcano plot showing association of protein abundance and co-clustering mutation in the same protein using the CPTAC retrospective and prospective cohorts of breast (BR), ovarian (OV), and colorectal (CO) cancers.

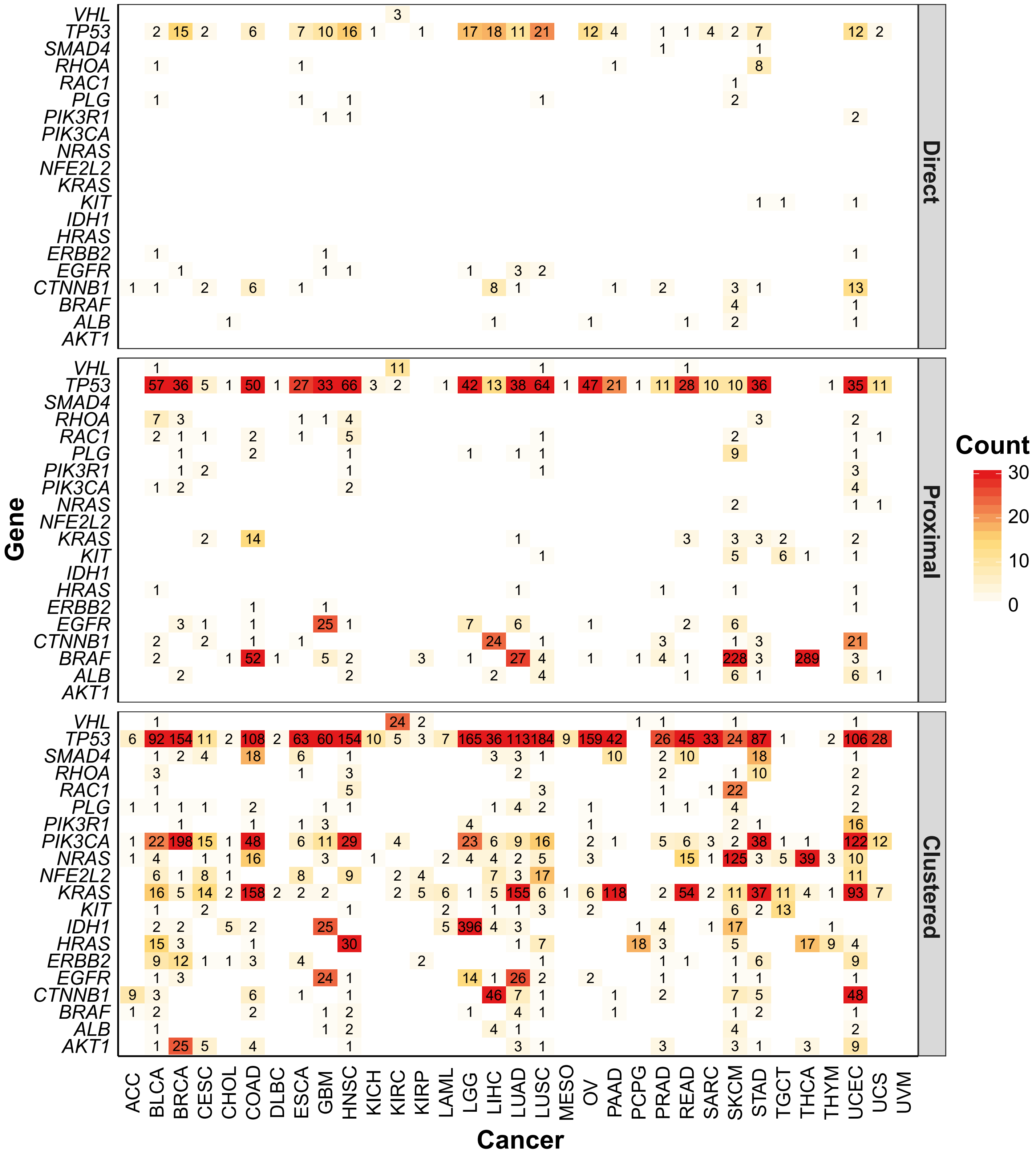


**Supplementary Figure 7. Count of missense mutations** (1) directly overlapping phosphosites, (2) proximal to phosphosites, and (3) co-clustering with phosphosites across 33 cancer types.

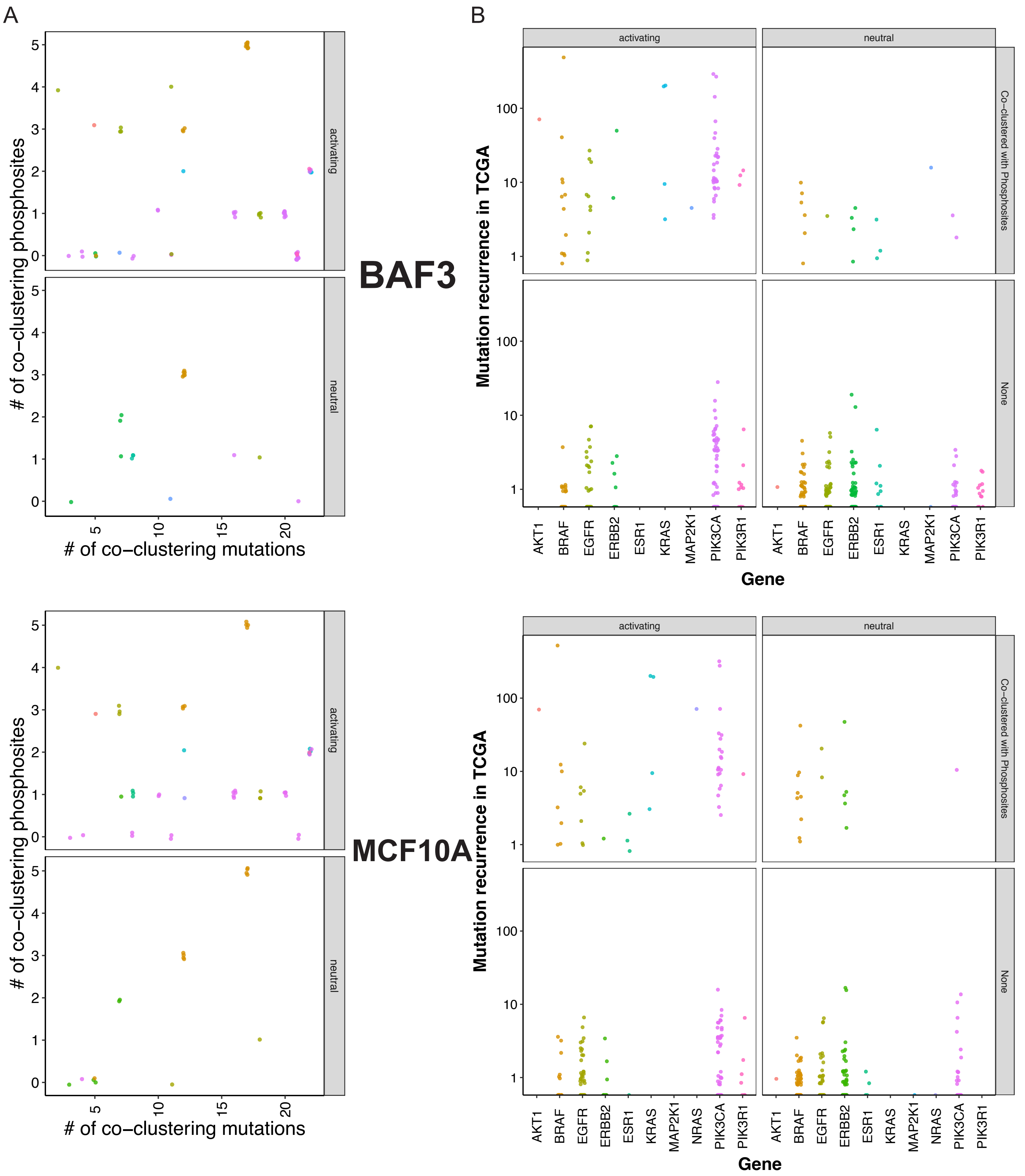


**Supplementary Figure 8. The relationship between mutation functionality versus co-clustering mutation counts, mutation recurrence, and co-clustering with phosphosites. (A)** Mutation functionality versus the counts of co-clustering phosphosites (y-axis) and mutations (x-axis). **(B)** Mutation functionality versus the mutation recurrence in TCGA by tested genes. For both (A) and (B), each dot represented one tested mutation, and the activating vs. neutral functionality labels were defined by results in BAF3 (top) and MCF10A (bottom)

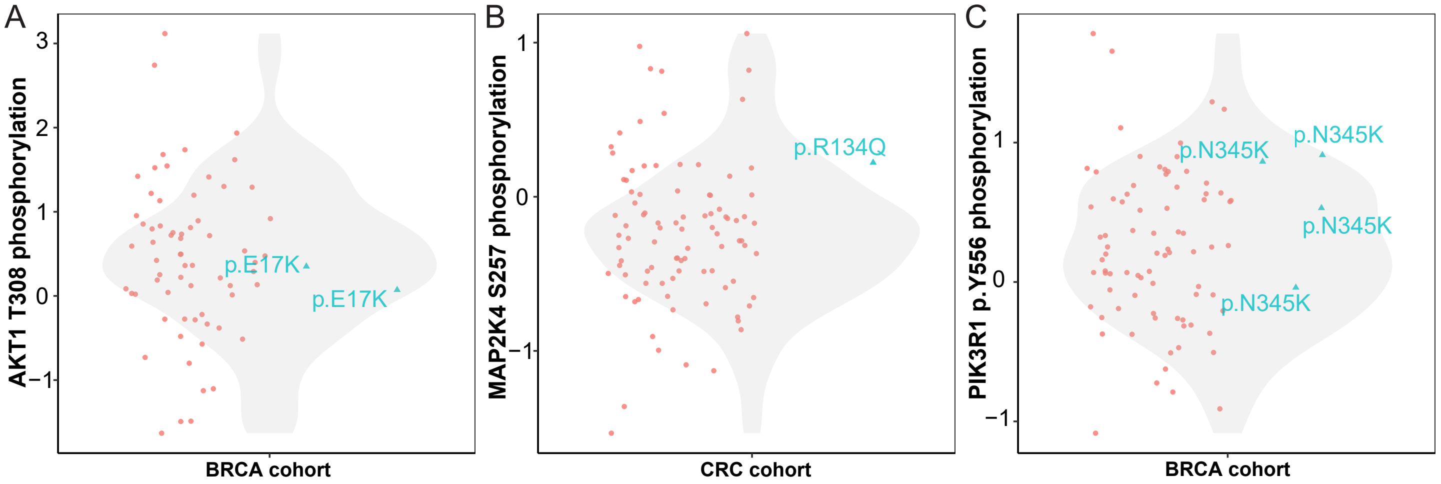


**Supplementary Figure 9. Potential interaction between co-clustering phosphosites and mutations in the same tumor sample.** (A) The two breast cancer samples carrying AKT1 p.E17K do not show differential phosphorylation of the co-clustering phosphosite p.T308. (B) The colorectal cancer sample carrying MAP2K4 p.R134Q mutation and its co-clustering phosphosite p.S257 level. (C) The 4 breast cancer samples carrying PIK3R1 p.N345K mutation and its co-clustering phosphosite p.Y556 levels.
